# Changes to the mtDNA copy number during yeast culture growth

**DOI:** 10.1101/2021.09.02.458779

**Authors:** Ben Galeota-Sprung, Amy Fernandez, Paul Sniegowski

## Abstract

We show that the mtDNA copy number in growing cultures of the yeast *Saccharomyces cerevisiae* increases by a factor of up to 4, being lowest (∼10 per haploid genome) and stable during rapid fermentative growth, and highest at the end of the respiratory phase. When yeast are grown on glucose, the onset of the mtDNA copy number increase coincides with the early stages of the diauxic shift, and the increase continues through respiration. A lesser yet still substantial copy number increase occurs when yeast are grown on a nonfermentable carbon source, i.e. when there is no diauxic shift. The mtDNA copy number increase during and for some time after the diauxic shift is not driven by an increase in cell size. The copy number increase occurs in both haploid and diploid strains, but is markedly attenuated in a diploid wild isolate that is a ready sporulator. Strain-to-strain differences in mtDNA copy number are least apparent in fermentation and most apparent in late respiration or stationary phase. While changes in mitochondrial morphology and function were previously known to accompany changes in physiological state, it had not been previously shown that the mtDNA copy number changes substantially over time in a clonal growing culture. The mtDNA copy number in yeast is therefore a highly dynamic phenotype.

## Introduction

Mitochondria are membrane-enclosed organelles that originate from an ancient symbiosis (Gray, 2017; Sagan, 1967). Mitochondria enable a variety of capabilities, the most important of which is the production of ATP via aerobic respiration. To a varying degree across the eukaryotic tree of life, most originally mitochondrial genes have been assimilated into the nuclear genome (Malina et al., 2018), but in nearly all cases an independent mitochondrial genome (mtDNA) is maintained within the mitochondria. Generally there are multiple copies of mtDNA per individual cell. In multicellular organisms the copy number may vary substantially. For example, in humans, there are hundreds to thousands to tens of thousands of mtDNA copies per cell with considerable variation between tissue type (D’Erchia et al., 2015; Dimmock et al., 2010; Miller et al., 2003; Wai et al., 2010) as well as substantial inter-individual variation (Kaaman et al., 2007). In humans, deficiency in mtDNA copy number is associated with a variety of human disease phenotypes (Elpeleg et al., 2002; Kornblum et al., 2013; Pyle et al., 2016; Zeviani & Di Donato, 2004).

The yeast *Saccharomyces cerevisiae* is an important model system for studying various aspects of mitochondrial biology (Shadel, 1999). The mitochondrial genome of *S. cerevisiae* encodes only 8 major proteins (the vast majority of the ∼1000 mitochondrial proteins are encoded in the nuclear genome), of which 7 are subunits of various respiratory enzyme complexes of the inner membrane (Malina et al., 2018). The mtDNA copy number varies somewhat by strain (De Chiara et al., 2020). An early study (Hall et al., 1976) found that 16-25% of all cellular DNA is mitochondrial; given an mtDNA genome of 75-85 kbp and a nuclear genome of ∼12 Mbp, this suggests a copy number of ∼25 to ∼50 per haploid genome, a range generally borne out by subsequent studies (Lebedeva & Shadel, 2007; Solieri, 2010; Williamson, 2002), though copy numbers as high as 200 have been observed in diploids (Miyakawa et al., 2004). Two recent studies have found ∼18-20 mtDNA copies per haploid cell, and up to roughly threefold variability introduced by various gene deletions (Göke et al., 2019; Puddu et al., 2019).

In yeast, as in other organisms, mtDNA is packed into protein-DNA complexes call nucleoids (Miyakawa, 2017; Westermann, 2014; Williamson, 2002; Williamson & Slonimski, 1976). These contain mostly linear mtDNA in polydisperse (not in units of discrete genomes) linear tandem arrays containing on average one to two copies of the mitochondrial genome (Freel et al., 2015; Maleszka et al., 1991), though under anaerobic conditions there may be many fewer nucleoids with many more mtDNA per nucleoid (Shiiba et al., 1997). The mitochondria within the yeast cell form a highly dynamic network in which fusion and fission events occur very frequently (Jakobs et al., 2003; Simon et al., 1997). There is generally a single “giant” mitochondrion that is much bigger than the others, and the number of spatially distinct mitochondria varies with physiological state, being lowest in rapidly growing cells and highest in stationary phase cells (Stevens, 1981).

*S. cerevisiae* that are grown with glucose as a carbon source characteristically ferment it into ethanol and then transition into a fully respiring physiology—the diauxic shift—upon the exhaustion of glucose supplies. The diauxic shifts initiates a substantial reorganization of yeast metabolism in which expression patterns for different sets of proteins (e.g. those associated with oxidative phosphorylation, stress response, and the glyoxylate cycle) change dramatically in a coordinated and staged manner (DeRisi et al., 1997; J. P. Murphy et al., 2015; Zampar et al., 2013). Much of this metabolic reorganization occurs at the mitochondrial level as the mitochondrial role transitions from biosynthetic hub to energy generator during the diauxic shift, which should probably be considered a distinct metabolic phase (Bartolomeo et al., 2020).

In this study, we investigate changes in mtDNA copy number over time by sampling repeatedly from growing cultures of *S. cerevisiae*, using haploid and diploid variants of both a common laboratory strain and a natural isolate. We find that the mtDNA copy number during fermentation is surprisingly low (∼10 per haploid genome). The onset of the diauxic shift is associated with the start of a large mtDNA copy number increase that, from rapid fermentative growth through to stationary phase, may be as high as 4-fold or more. This suggests that the metabolic reorganization of the diauxic shift is associated with a substantial increase in mtDNA copy number. There is also a smaller mtDNA increase even when there is no diauxic shift (that is, when there is no fermentation phase). We confirm the reverse phenomenon of the copy number rise: when slowly growing cells are transferred to fresh media, the mtDNA copy number drops rapidly. We find that differences between the strains are most apparent after the fermentation phase. We find linear effects of ploidy, and find that the copy number increase has a complex relationship with cell size. These results suggest that the mtDNA copy number is a highly dynamic and plastic phenotype in yeast, and point to new directions for research into the regulation of mtDNA copy number in yeast.

## Results

### The mtDNA copy number increases over time in growing cultures

In our first experiment, we grew replicate cultures in rich liquid media supplemented with glucose (YPD). We employed haploid and diploid strains constructed from both a standard laboratory background (W303) and a natural woodland isolate. For all cultures, the first DNA extraction was performed at an absorbance corresponding to rapid fermentative growth (OD600=0.5), with subsequent extractions taking place at ∼24-hour intervals. The relative ratio of mtDNA to nuclear DNA (mt/nDNA) was assayed by qPCR. We found (Figs 1A, 1B) that the mt/nDNA ratio increased ∼3-fold for wild haploids, and over 4-fold for both haploid and diploid lab strains. The mt/nDNA ratio per haploid genome was similar across the two different ploidies, consistent with prior results (De Chiara et al., 2020). The exception was wild diploids, for which mtDNA copy number first doubled but subsequently declined, likely because of their strong propensity to sporulate spontaneously under these conditions, while laboratory strains tend to sporulate only if specifically induced.

**Fig. 1.**
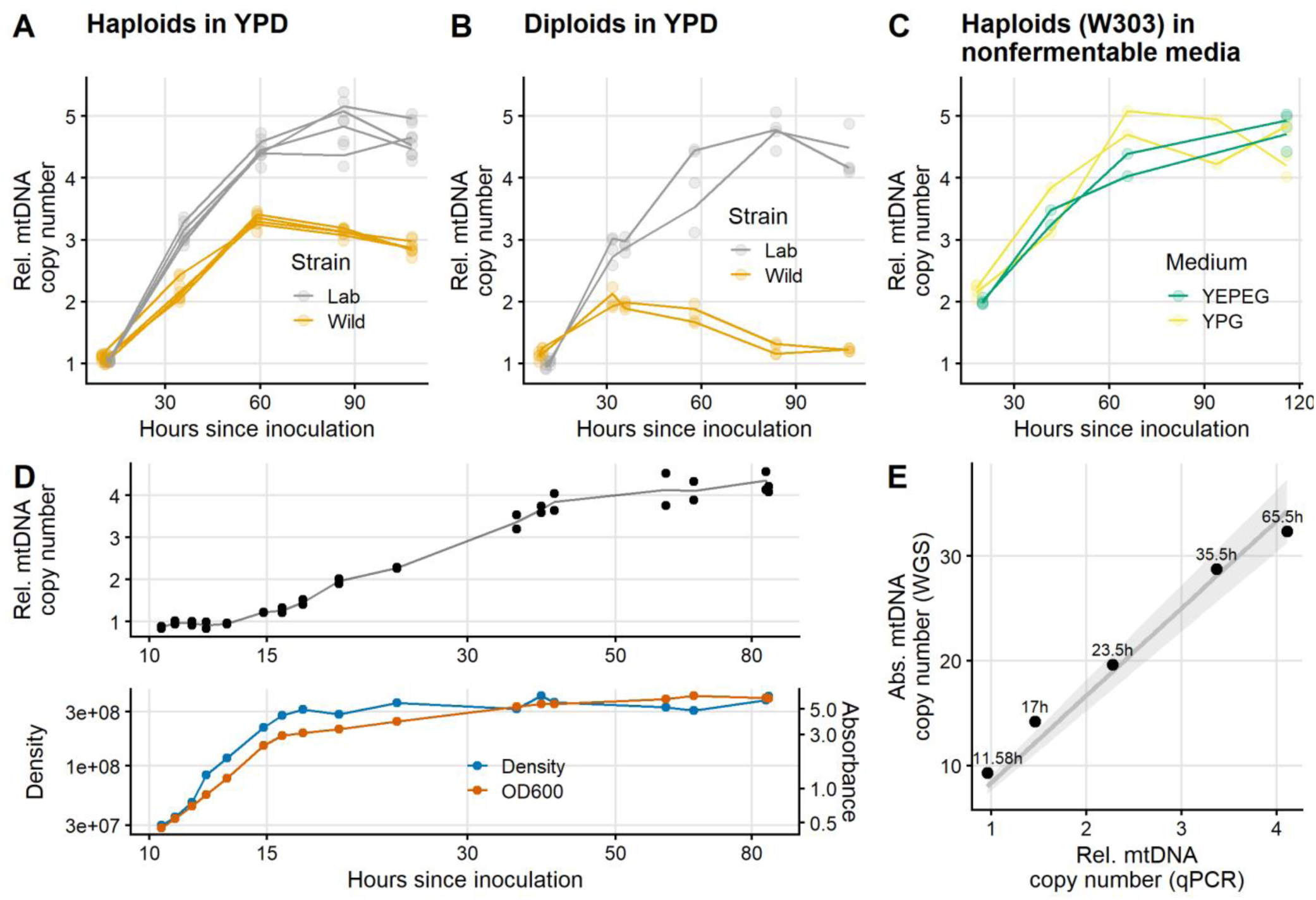
Relative mtDNA copy number over time in replicate cultures grown in rich media supplemented with glucose (**A**, haploids and **B**, diploids), or glycerol and glycerol + ethanol (**C**, W303 haploids only). DNA extractions were carried out for each culture at approximately 24-hour intervals. Two replicate qPCR measurements were performed for each DNA extraction; the lines track the mean of the two measurements. The first extraction was always performed at OD600 = 0.5 (∼11h after inoculation in YPD) which corresponded to ∼3e07 cells/mL in haploid strains and ∼2.4e07 cells/mL in diploid strains. **D:** Detailed mtDNA copy number dynamics of a single haploid W303 culture grown in YPD, with time shown on a log scale to better visualize the earlier data points. For 5 of 17 timepoints, the measurement of mtDNA copy number by qPCR was validated by a whole genome-sequencing (WGS) read-depth assay (**E**).

### mtDNA copy number estimation by WGS

Having established the general fact of mtDNA copy number increase for a variety of strains and conditions, we focused on the haploid lab strain for further study. We next sampled more extensively from a single culture grown on glucose, assaying the mt/nDNA ratio by qPCR at a total of 17 timepoints. These results demonstrated that the mt/nDNA ratio was stable during fermentation prior to the slowdown of growth associated with glucose exhaustion (Fig. 1D, early time points). As a control to rule out any effect of total time in culture, we also confirmed that the relative mtDNA copy number remains at ∼1 when a culture is repeatedly transferred while still in fermentation for over 60 hours (Fig. S6).

For 5 of those same 17 timepoints for which qPCR assays were performed, we carried out a WGS read-depth assay, the purpose of which was to verify the qPCR results as well as to estimate the absolute (rather than relative) mtDNA copy number. We found that the mtDNA copy number during fermentative growth for haploids is ∼9, eventually increasing to over 30, and that there is good agreement (Fig. 1E) between the qPCR and WGS assays as to the scale of the increase in copy number, though the WGS assay shows a slightly smaller fold increase (Fig. S1). As controls for the WGS assay, we assayed the copy number of various other repetitive genomic elements, which did not change over time (Fig. S2). We also found good agreement between short- and long-read WGS-based assays for mtDNA copy number (Figs. S3-S5).

### mtDNA copy number increase and the diauxic shift

*S. cerevisiae* is a Crabtree-positive yeast: it ferments glucose into ethanol even when oxygen is present, and then approaching glucose exhaustion enters the diauxic shift, transitioning from fermentation of glucose to oxidative respiration of ethanol. We were interested in the association of the diauxic shift with the observed rise in mtDNA copy number. Therefore, we conducted another dense-sampling experiment in which ethanol and glucose concentrations were regularly measured. We also investigated changes in cell size in this experiment.

We found (Fig. 2) that the increase in mtDNA copy number increase begins at the very onset of the diauxic shift, as the rate of cellular proliferation slows and prior to total glucose exhaustion. The mtDNA copy number increase also after the diauxic shift and appears to last throughout respiration. The rate of mtDNA copy number increase was faster during the diauxic shift and slower during respiratory phase, though the difference in rate (0.15 hr^-1^ vs 0.09 hr^-1^) is not statistically significant in our data (Fig. S8).

**Fig. 2.**
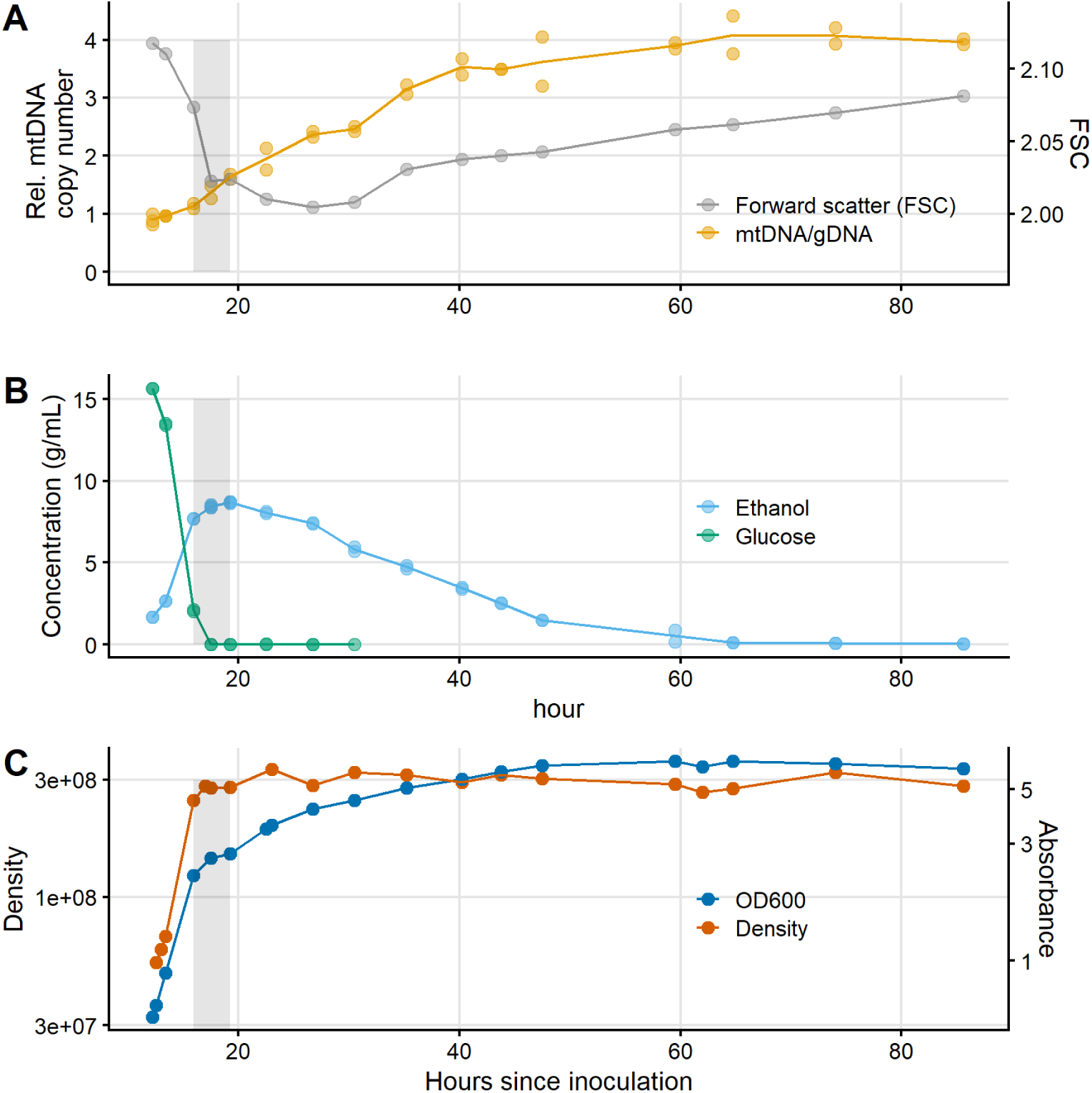
For a haploid W303 culture grown in YPD, repeated estimate of (**A**) mtDNA copy number and cell size, (**B**) ethanol and glucose concentration, and (**C**) absorbance and density. The gray box shows the approximate boundaries of the diauxic shift.

The portion of the increase in mtDNA copy number that occurs during respiratory phase does not seem to require the diauxic shift to have previously occurred: we observed that cultures grown on a nonfermentable carbon source—glycerol alone or glycerol + ethanol—also underwent an mtDNA copy number increase (Fig. 1C). This is increase is about half the total magnitude of the increase that occurred during growth on glucose, as the first measurement of mtDNA copy number (conducted at OD600=0.5, as in the glucose cultures) is about twice as high as during fermentative growth and the final measurements are indistinguishable between the two conditions.

### mtDNA copy number changes and cell size

We estimated cell size by flow cytometry (Fig. 2A), supplemented with imaging via light microscopy (Fig. S7). We observed a decline in cell size that carried through and for some time after the diauxic shift before a later increase in cell size in respiratory phase. This decline in cell volume is consistent with other observations in whole (Brauer et al., 2005) or in part (Bartolomeo et al., 2020). Slightly more than half of the mtDNA copy increase occurred while cell size was declining. Thus there is an initially negative association between rate of change of cell size and rate of change of mtDNA copy number that eventually shifts to a positive association. It should be noted that this negative rate relationship does not preclude the possibility that at any given instant there is always a positive relationship within the population between cell size and mtDNA copy number.

### Decrease in mtDNA copy number upon transfer from stationary phase

Given that the mtDNA copy number increases during growth from dilute to saturated culture, we reasoned that the reverse process must take place: a decline in mtDNA copy number accompanies the resumption of growth that occurs upon transfer from a saturated culture to fresh media (Williamson & Moustacchi, 1971). In order to observe this process we transferred a relatively large number of late respiratory phase cells, sufficient that extractions could be immediately performed, to fresh YPD and began regular extractions immediately. After a lag until the resumption of cellular proliferation of about an hour, the expected drop in mtDNA copy number occurred rapidly (Fig. 3) over the next 5 hours.

**Fig. 3.**
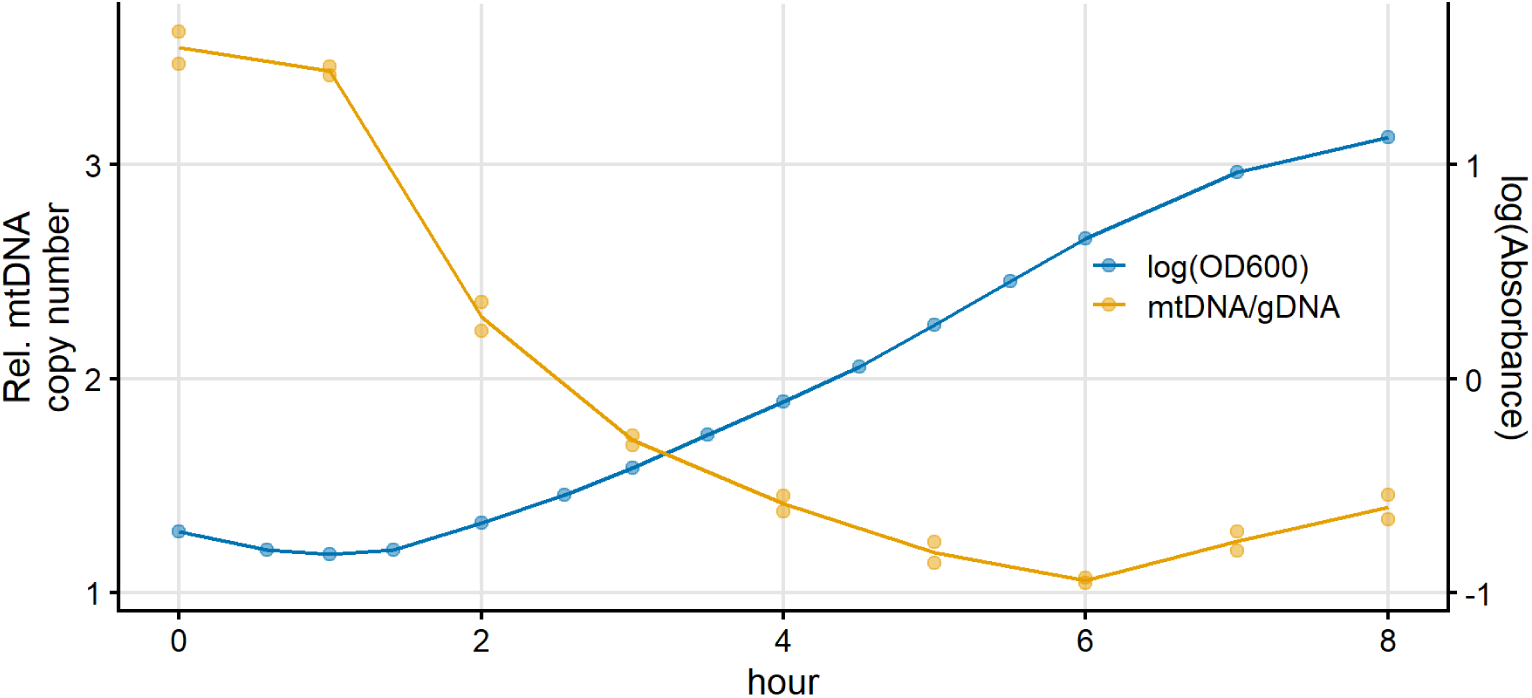
The drop in copy number immediately after transfer to fresh YPD. A saturated culture of the haploid lab strain was transferred, diluting 1/30, to fresh YPD. The relatively mild dilution permitted DNA extractions of small portions of the culture to begin immediately after transfer. Extractions for qPCR assays, and density and absorbance measurements, were performed at 1-hour intervals for 8 hours. From hours 1-3 the mtDNA copy number falls be a factor of 2 and the population grows by a factor of 2, suggesting that there was no mtDNA production during this time.

The initial magnitude of the decline in mtDNA copy number compared to the magnitude of the increase in population suggests that virtually no mtDNA production at all was taking place from hours 1-3, during which the density increased and the mtDNA copy number decreased by a factor of ∼2. Averaged over hours 1-6, the difference in Malthusian growth parameter between population growth rate and the mtDNA production rate is 0.25 hour^-1^.

## Discussion

The main result of this paper is that the mtDNA copy number rises dramatically during the course of typical batch culture growth on glucose, by a factor of up to ∼4-fold, depending on strain. The copy number is lowest during rapid fermentative growth and increases during and after the switch to respiration. While it was previously thought that the copy number does not change very much over different growth conditions (Chen & Butow, 2005), some degree of copy number change is not necessarily surprising. It has long been noted that the morphology of the yeast mitochondrial network is plastic and undergoes dramatic changes in response to different environmental conditions (Miyakawa, 2017), and one previous study (Miyakawa et al., 2004) reported a ∼40% difference in mtDNA copy number between growth in aerobic and anaerobic conditions. The scale of the mtDNA copy number change that we report here, however, has not been previously reported.

We observe that the mtDNA copy number is stable during fermentation as long as glucose is abundant (Fig. 1D). The yeast transcriptome during fermentation is a mostly steady state (Pelechano & Pérez-Ortín, 2010) until the diauxic shift nears; thus it appears that this period of transcriptome stability is associated with copy number stability. While we find a fairly large strain-to-strain difference in mtDNA copy number (Fig. 1), this difference is minimized during fermentation, perhaps suggesting relatively tight control of a minimum complement of mitochondria during fermentation. We find that the absolute copy number, as measured by WGS, is 9 copies per haploid genome (Fig. 1E) during rapid fermentative growth. This is a lower copy number than is usually reported, perhaps because the copy number has not typically been assayed during rapid fermentative growth. We find that the absolute copy number varies over culture conditions to reach ∼30-40 in haploids and just about twice that in diploids, a linear scaling with ploidy that has previously been reported (De Chiara et al., 2020, Grimes et al., 1974, Puddu et al. 2019, though see Göke et al 2020 for a counterexample).

The increase in mtDNA copy number begins before glucose has been fully exhausted (Fig. 2). This timing is very similar to the timing of the coordinated induction of stress and oxidative phosphorylation proteins that marks the earliest onset of the diauxic shift prior to complete glucose depletion (DeRisi et al., 1997; J. P. Murphy et al., 2015; Zampar et al., 2013). All but one of the 8 protein-coding genes on the *S. cerevisiae* mitochondrial genome are subunits of an oxidative phosphorylation enzyme complex. Thus it would seem that the onset of the mtDNA copy number increase coincides with the onset of increased expression of most mitochondrially located genes. Data from Murphy et al (2015) that illustrates this point is re-plotted in Fig. S9. The causality that determines this temporal correlation is open to further investigation.

Although the mtDNA copy number increase is fairly rapid once it begins (Fig. S8), it is important to point out that our data do not suggest an absolute (i.e., per unit of time) increase in the rate of mtDNA replication upon the diauxic shift, because the rate of cellular reproduction slows dramatically at this time. In fact, our data suggest that the rate of mtDNA production decreases at the diauxic shift, but that it decreases less than the rate of cellular reproduction does, so that there is a net increase in mtDNA production relative to each cell cycle. This is possible because in *S. cerevisiae*, mtDNA replication is decoupled from the cell cycle (Williamson & Moustacchi, 1971). We illustrate the point that relative rates of nuclear and mitochondrial genome growth determine the copy number with the toy model depicted in Fig. S10, in which staggered alternation of equal genomic and mitochondrial growth rates leads to a cyclical rise and fall in the copy number. Indeed, we observe that the mtDNA copy number drops sharply upon transfer to fresh media (Fig. 3): our data suggests that in such conditions, the lag for mitochondrial production to resume is longer than the lag for cellular reproduction to resume, causing a very rapid drop in mtDNA copy number until mtDNA production restarts and subsequently catches up to the budding rate.

The timing of the onset of the mtDNA copy number increase observed here also corresponds well with the mitochondrial volume increase reported by Bartolomeo et al (2020). This increase in the absolute mitochondrial volume per cell begins early in the diauxic shift, just as the copy number increase does, and there is an initial doubling in mitochondrial volume in roughly the same timeframe as we observe a doubling in mtDNA copy number. The implication is that absolute mitochondrial volume per cell is well-correlated with mtDNA copy number per cell, though further work must be done to establish this.

The relationship between mtDNA copy number and cell size is more complex. The number of mtDNA nucleoids is correlated with the length of the mitochondrial network (Osman et al., 2015), which itself is correlated with cell volume (Rafelski et al., 2012). This suggests that the mtDNA copy number is correlated with cell volume, with the caveat that the number of nucleoids and the mtDNA copy number is not necessarily correlated. Indeed, at a single point in time during fermentation, the diauxic shift, or respiration, cell volume and mtDNA copy number are well-correlated within the population, as shown by Bartolomeo et al. (2020). However, we find that cell volume actually decreases after the diauxic shift, consistent with reports by Bartolomeo et al. (2020) and Braur et al. (2005). Roughly speaking, the first half of the mtDNA copy number increase occurs while cell size is decreasing (Fig. 2A), so that there is a negative correlation between the rate of change in copy number and the rate of change in cell volume. Then, in late respiration, cell volume increases while the mtDNA copy number is still increasing, and the sign of the correlation become positive.

We observe a copy number increase during respiration even when there was never any fermentation (Fig. 1C). This suggests the possibility that there may be two separable phenomena: an approximate doubling of mtDNA copy number with the diauxic shift, and then a further rise in respiration that occurs whether or not there was a diauxic shift. To the extent that either of these phenomena are adaptive, which is not necessarily the case, we might hypothesize that the initial and more rapid copy number increase during the diauxic shift reflects a demand for more mitochondria as their role transitions from fatty acid production to primary energy generator, while the slower copy number increase that extends throughout respiration to stationary phase might reflect preparation for the sexual mode of reproduction of yeast, that is, sporulation, which occurs in conditions of nitrogen starvation without a fermentable carbon source as well as under other types of nutrient deprivation (Freese et al., 1982). During sporulation, the single genome doubling of meiosis is not accompanied by a corresponding increase in mtDNA copy number, and only a bit more than half of the total mtDNA of the parent diploid is included within one of the four haploid spores (Brewer & Fangman, 1980). Thus a sporulating diploid might require a high mtDNA copy number in order for each of its four haploid offspring to possess an adequate number of mtDNA copies. Indeed, the copy number dynamics of the wild diploid strain (Fig. 1B) are very different from the other strains. Unlike laboratory diploid strains, this strain is a very ready sporulator and we observed that the great majority of cells had sporulated by the end of the experiment, as has been previously reported for some wild isolates grown on YPD (H. A. Murphy & Zeyl, 2010). In this strain only, the mtDNA copy number falls after the initial doubling during the diauxic shift. Little remains known about the natural ecology of *S. cerevisiae* and its propensity to sporulate in natural environments (Knight & Goddard, 2016).

## Conclusions

In summary, we find that the mtDNA copy number of *S. cerevisiae* is quite low (∼10 per haploid genome) during the rapid growth of fermentation, and increases substantially during the diauxic shift and respiratory phases. An implication of this result is that future studies of the genetic control of mtDNA copy number should take care to compare copy number at similar culture phases, in order to filter out apparent variation in copy number that might be caused by variation in growth dynamics. The wholesale reorganization of yeast metabolism that begins with the onset of the diauxic shift may be initiated from the mitochondria (Bartolomeo et al., 2020). We observe that the copy number increase begins very early during the diauxic shift, when both transcriptional activity and mitochondrial volume are known to begin to change rapidly. The causal connection between these changes and the mtDNA copy number change is open to further study.

## Methods

### Strains

yJHK112, a haploid, prototrophic, heterothallic, *MATa, BUD4*-corrected, and ymCherry-labeled W303 strain (Koschwanez et al., 2013) was used as the haploid lab strain in this paper. We created the diploid lab strain from this strain by transforming (Gietz and Schiestl 2007) the haploid lab strain with plasmid pRY003, temporarily providing a functional *HO* locus allowing mating type switching and subsequent mating. pRY003 was a gift from John McCusker (Addgene plasmid #81043; http://n2t.net/addgene:81043; RRID:Addgene_81043). The ploidy of the resulting strain was confirmed by (1) ability to produce tetrads after plating to sporulation media; (2) by flow cytometry for total genomic content; and (3) by the presence of a PCR product for both the *MATa* and *MATα* loci. YPS623, the wild diploid strain used in this paper, was isolated from Mettler’s Woods, NJ following procedures described in (Sniegowski et al., 2002). YPS 2066, the haploid wild strain used in this paper, originated from an ascospore of YPS623, isolated by microdissection, and subsequently made heterothallic by knocking out the *HO* locus by standard genetic techniques.

### Growth conditions

Growth was carried out in liquid media in Erlenmeyer flasks kept at 30 C and shaken at 200 rpm. For all experiments, an initial growth cycle was carried out in 6 mL media in a 50 mL flask upon revival from frozen storage. Unless otherwise noted, all experiments described in this paper were carried out in 50 mL media in a 250 mL flask. Except for the experiment described in Fig. 3, all transfers to start experiments were 1:2000 (25 ul into 50 mL) dilutions from pregrown cultures. Three kinds of media were used: YPD (also known as YEPD) (2% peptone, 2% glucose, 1% yeast extract); YPG (2% peptone, 3% glycerol, 1% yeast extract); and YPEG (2% peptone, 2.6% glycerol, 2.6% ethanol, 1% yeast extract).

### Measurement of culture properties

Density measurements were performed by manually counting the number of cells, via microscope, in an improved Neubauer-style hemocytometer. Depending on culture density, dilutions of up to 1/200 (in water) were performed before counting. Still-attached buds were counted as a separate cell if having a diameter greater than half that of the mother.

Absorbance was measured at 600 nm (OD600) by standard methods. If the OD600 was greater than 1, the culture was diluted 1/10 in the appropriate media to in order to obtain a reading within the better part of the dynamic range of the spectrophotometer. Using a data set of dual (diluted and undiluted) measurements to interpolate, all diluted OD600 measurements were converted to the scale of undiluted measurements so that only one scale of measurement is presented.

To measure glucose and ethanol concentrations, 0.5 mL culture samples were spun down and the supernatant stored at -20 C. Samples were then defrosted for glucose and ethanol measurement. R-Biopharm kit 10 176 290 035 and Megazyme kit K-GLUHK-110A were used to measure ethanol and glucose concentration, respectively. Both kits were used according to the manufacturer’s instructions.

### DNA extractions

DNA extractions were performed using Zymo Quick-DNA Fungal/Bacterial kits (D6005), which employ glass beads to physically lyse cells followed by a spin column-based purification. In order to limit any effects of different extraction conditions, and because extractions were performed from growing cultures at different time points and hence different densities, within a single experiment the extraction volume was adjusted so that the absolute number of cells extracted was similar for each timepoint, As an exception to this practice, extractions destined for whole-genome sequencing were larger in cell number than extractions destined for qPCR only.

In early exploratory experiments, similar results as to those described here were obtained with both purely enzymatic extraction protocols (Zymo D2002) as well as a non-kit-based glass beads/phenol:chloroform extraction protocol.

### qPCR

qPCR assays for mtDNA content were performed using the relative standard curve method. For both lab and wild strains, a single large-scale extraction from the haploid strain was performed when the culture was in fermentation, at absorbance OD600=0.6. For every qPCR run, a fivefold serial dilution of this single extract served as the standard curve. Each sample was run with two sets of primers, identical to the primers employed in Puddu et al. (2019). We employed one set of primers to target a region of nuclear DNA and another to amplify a region of mitochondrial DNA. The nDNA primer set (fwd: TGCTTTGTCAAATGGATCATATGG, rev: CCTGGAACCAAGTGAACAGTACAA) targeted a short region of GAL1 (chr II: 280382–280459) and the mtDNA primer set (fwd: CACCACTAATTGAAAACCTGTCTG, rev: GATTTATCGTATGCTCATTTCCAA) targeted a short region of COX1 in the mtDNA (25574– 25686). For each sample, the quantity of mtDNA and nDNA relative to the standard curve was computed, and the ratio of the two recorded as the relative mtDNA copy number. Threshold calculations, curve fitting, and quantity calculations were performed by the QuantStudio 3 software. All wells were run in triplicate per run and outliers from each triplet were occasionally removed manually. Each qPCR run was performed on a 96-well plate allowing a maximum of 11 unknown samples to be assayed for relative mtDNA copy number (11 samples x 2 primer sets x in triplicate = 66 wells, plus 5 standards x 2 primer sets x in triplicate = 30 wells, for a total of 96 wells).

### Sequencing and copy number analysis

The mtDNA copy number analysis was conducted similarly to Puddu et al. (2019) and was carried out working from short-read (Illumina) WGS data as follows (also see `sequencing_pipeline.txt` in the online data). Reads were first processed by `trim_galore` with options `--nextera --stringency 3 --paired --quality 20 --length 50`. Reads were mapped to the S288C R64 reference genome by bwa. Duplicate reads were removed by `biobambam2’s bammarkduplicates` and then read depth computed by `samtools depth`. The mtDNA genome includes many regions of high AT content that are not covered well; hence the mtDNA copy number was computed as the median read depth in a region of the mtDNA genome with regular coverage (COX1 from 14000-20000; see Fig. S3) divided by the median read depth in the nuclear genome. mtDNA copy number calculated using COX3 instead (79213– 80022) was very similar. As controls, we calculated copy numbers for other repetitive elements (Fig. S2), which did not change over the course of culture growth.

Data presented in Figs. 1E and S1 was produced from short-read sequencing. Initially, we suspected that the coverage from short-read sequencing would be too irregular to permit accurate copy number estimates. Therefore, in preliminary experiments, we compared short-read (Illumina) and long-read (Nanopore) sequencing from the same samples. While the coverage for the Nanopore data was indeed somewhat smoother (Figs. S3 and S4), the mitochondrial coverage for the short-read data was acceptably regular. Indeed, the two methods gave near-identical results (Table S5).

### Cell size

We used forward scatter (FSC) as a proxy for cell size (Zakhartsev & Reuss, 2018). Culture samples of varying volume (to account for changing culture density) were fixed in 70% ethanol and stored at 4C, then spun down, washed once, and resuspended in sodium citrate buffer. After a 1/10 dilution, samples were sonicated twice for 10s at 30% power, examined microscopically to ensure sonication was effective, and then diluted 1/20 for flow cytometry on a Guava EasyCyte. Samples were run for 90s at a flow rate of Very Low, with doublet discrimination performed by removing off-diagonal points on FSC-W vs FSC-H.

We obtained an independent set of measurements of cell size by light microscopy. Live cells were imaged on a Nexcelom Cellometer X2 and processed using an object detection and measurement pipeline (available in the online supplement) implemented in CellProfiler (McQuin et al., 2018). In this pipeline, all processed images were reviewed and any cell aggregations that were not declustered properly were manually removed from the output (sample images included in Fig. S7).

## Data and code

All code to generate figures in the submitted manuscript, along with the underlying data, is available for review here: https://datadryad.org/stash/share/EVqfjBsUcDtCQ_9T7_H8LGkXJntOeBzoZ0X8Er86gl0

## References

Bartolomeo, F. D., Malina, C., Campbell, K., Mormino, M., Fuchs, J., Vorontsov, E., Gustafsson, C. M., & Nielsen, J. (2020). Absolute yeast mitochondrial proteome quantification reveals trade-off between biosynthesis and energy generation during diauxic shift. Proceedings of the National Academy of Sciences, 117(13), 7524–7535. https://doi.org/10.1073/pnas.1918216117

Brauer, M. J., Saldanha, A. J., Dolinski, K., & Botstein, D. (2005). Homeostatic Adjustment and Metabolic Remodeling in Glucose-limited Yeast Cultures. Molecular Biology of the Cell, 16(5), 2503–2517. https://doi.org/10.1091/mbc.e04-11-0968

Brewer, B. J., & Fangman, W. L. (1980). Preferential inclusion of extrachromosomal genetic elements in yeast meiotic spores. Proceedings of the National Academy of Sciences, 77(9), 5380–5384. https://doi.org/10.1073/pnas.77.9.5380

Chen, X. J., & Butow, R. A. (2005). The organization and inheritance of the mitochondrial genome. Nature Reviews Genetics, 6(11), 815–825. https://doi.org/10.1038/nrg1708

De Chiara, M., Friedrich, A., Barré, B., Breitenbach, M., Schacherer, J., & Liti, G. (2020). Discordant evolution of mitochondrial and nuclear yeast genomes at population level. BMC Biology, 18(1), 49. https://doi.org/10.1186/s12915-020-00786-4

D’Erchia, A. M., Atlante, A., Gadaleta, G., Pavesi, G., Chiara, M., De Virgilio, C., Manzari, C., Mastropasqua, F., Prazzoli, G. M., Picardi, E., Gissi, C., Horner, D., Reyes, A., Sbisà, E., Tullo, A., & Pesole, G. (2015). Tissue-specific mtDNA abundance from exome data and its correlation with mitochondrial transcription, mass and respiratory activity. Mitochondrion, 20, 13–21. https://doi.org/10.1016/j.mito.2014.10.005

DeRisi, J. L., Iyer, V. R., & Brown, P. O. (1997). Exploring the Metabolic and Genetic Control of Gene Expression on a Genomic Scale. Science. https://doi.org/10.1126/science.278.5338.680

Dimmock, D., Tang, L.-Y., Schmitt, E. S., & Wong, L.-J. C. (2010). Quantitative Evaluation of the Mitochondrial DNA Depletion Syndrome. Clinical Chemistry, 56(7), 1119–1127. https://doi.org/10.1373/clinchem.2009.141549

Elpeleg, O., Mandel, H., & Saada, A. (2002). Depletion of the other genome-mitochondrial DNA depletion syndromes in humans. Journal of Molecular Medicine, 80(7), 389–396. https://doi.org/10.1007/s00109-002-0343-5

Freel, K. C., Friedrich, A., & Schacherer, J. (2015). Mitochondrial genome evolution in yeasts: An all-encompassing view. FEMS Yeast Research, 15(fov023). https://doi.org/10.1093/femsyr/fov023

Freese, E. B., Chu, M. I., & Freese, E. (1982). Initiation of Yeast Sporulation by Partial Carbon, Nitrogen, or Phosphate Deprivation. Journal of Bacteriology, 149(3), 840–851. https://doi.org/10.1128/jb.149.3.840-851.1982

Göke, A., Schrott, S., Mizrak, A., Belyy, V., Osman, C., & Walter, P. (2019). Mrx6 regulates mitochondrial DNA copy number in Saccharomyces cerevisiae by engaging the evolutionarily conserved Lon protease Pim1. Molecular Biology of the Cell, 31(7), 527–545. https://doi.org/10.1091/mbc.E19-08-0470

Gray, M. W. (2017). Lynn Margulis and the endosymbiont hypothesis: 50 years later. Molecular Biology of the Cell, 28(10), 1285–1287. https://doi.org/10.1091/mbc.e16-07-0509

Hall, R. M., Nagley, P., & Linnane, A. W. (1976). Biogenesis of mitochondria. XLII. Genetic analysis of the control of cellular mitochondrial DNA levels in Saccharomyces cerevisiae. Molecular & General Genetics: MGG, 145(2), 169–175. https://doi.org/10.1007/BF00269590

Jakobs, S., Martini, N., Schauss, A. C., Egner, A., Westermann, B., & Hell, S. W. (2003). Spatial and temporal dynamics of budding yeast mitochondria lacking the division component Fis1p. Journal of Cell Science, 116(10), 2005–2014. https://doi.org/10.1242/jcs.00423

Kaaman, M., Sparks, L. M., van Harmelen, V., Smith, S. R., Sjölin, E., Dahlman, I., & Arner, P. (2007). Strong association between mitochondrial DNA copy number and lipogenesis in human white adipose tissue. Diabetologia, 50(12), 2526–2533. https://doi.org/10.1007/s00125-007-0818-6

Knight, S. J., & Goddard, M. R. (2016). Sporulation in soil as an overwinter survival strategy in Saccharomyces cerevisiae. FEMS Yeast Research, 16(1), fov102. https://doi.org/10.1093/femsyr/fov102

Kornblum, C., Nicholls, T. J., Haack, T. B., Schöler, S., Peeva, V., Danhauser, K., Hallmann, K., Zsurka, G., Rorbach, J., Iuso, A., Wieland, T., Sciacco, M., Ronchi, D., Comi, G. P., Moggio, M., Quinzii, C. M., DiMauro, S., Calvo, S. E., Mootha, V. K., … Prokisch, H. (2013). Loss-of-function mutations in MGME1 impair mtDNA replication and cause multisystemic mitochondrial disease. Nature Genetics, 45(2), 214–219. https://doi.org/10.1038/ng.2501

Lebedeva, M. A., & Shadel, G. S. (2007). Cell Cycle- and Ribonucleotide Reductase-Driven Changes in mtDNA Copy Number Influence mtDNA Inheritance Without Compromising Mitochondrial Gene Expression. Cell Cycle, 6(16), 2048–2057. https://doi.org/10.4161/cc.6.16.4572

Maleszka, R., Skelly, P. J., & Clark-Walker, G. D. (1991). Rolling circle replication of DNA in yeast mitochondria. The EMBO Journal, 10(12), 3923–3929.

Malina, C., Larsson, C., & Nielsen, J. (2018). Yeast mitochondria: An overview of mitochondrial biology and the potential of mitochondrial systems biology. FEMS Yeast Research, 18(foy040). https://doi.org/10.1093/femsyr/foy040

McQuin, C., Goodman, A., Chernyshev, V., Kamentsky, L., Cimini, B. A., Karhohs, K. W., Doan, M., Ding, L., Rafelski, S. M., Thirstrup, D., Wiegraebe, W., Singh, S., Becker, T., Caicedo, J. C., & Carpenter, A. E. (2018). CellProfiler 3.0: Next-generation image processing for biology. PLOS Biology, 16(7), e2005970. https://doi.org/10.1371/journal.pbio.2005970

Miller, F. J., Rosenfeldt, F. L., Zhang, C., Linnane, A. W., & Nagley, P. (2003). Precise determination of mitochondrial DNA copy number in human skeletal and cardiac muscle by a PCR-based assay: Lack of change of copy number with age. Nucleic Acids Research, 31(11), e61–e61. https://doi.org/10.1093/nar/gng060

Miyakawa, I. (2017). Organization and dynamics of yeast mitochondrial nucleoids. Proceedings of the Japan Academy. Series B, Physical and Biological Sciences, 93(5), 339–359. https://doi.org/10.2183/pjab.93.021

Miyakawa, I., Miyamoto, M., Kuroiwa, T., & Sando, N. (2004). DNA Content of Individual Mitochondrial Nucleoids Varies Depending on the Culture Conditions of the Yeast Saccharomyces cerevisiae. Cytologia, 69(1), 101–107. https://doi.org/10.1508/cytologia.69.101

Murphy, H. A., & Zeyl, C. W. (2010). Yeast Sex: Surprisingly High Rates of Outcrossing between Asci. PLOS ONE, 5(5), e10461. https://doi.org/10.1371/journal.pone.0010461

Murphy, J. P., Stepanova, E., Everley, R. A., Paulo, J. A., & Gygi, S. P. (2015). Comprehensive Temporal Protein Dynamics during the Diauxic Shift in Saccharomyces cerevisiae. Molecular & Cellular Proteomics, 14(9), 2454–2465. https://doi.org/10.1074/mcp.M114.045849

Osman, C., Noriega, T. R., Okreglak, V., Fung, J. C., & Walter, P. (2015). Integrity of the yeast mitochondrial genome, but not its distribution and inheritance, relies on mitochondrial fission and fusion. Proceedings of the National Academy of Sciences, 112(9), E947–E956. https://doi.org/10.1073/pnas.1501737112

Pelechano, V., & Pérez-Ortín, J. E. (2010). There is a steady-state transcriptome in exponentially growing yeast cells. Yeast, 27(7), 413–422. https://doi.org/10.1002/yea.1768

Puddu, F., Herzog, M., Selivanova, A., Wang, S., Zhu, J., Klein-Lavi, S., Gordon, M., Meirman, R., Millan-Zambrano, G., Ayestaran, I., Salguero, I., Sharan, R., Li, R., Kupiec, M., & Jackson, S. P. (2019). Genome architecture and stability in the Saccharomyces cerevisiae knockout collection. Nature, 573(7774), 416–420. https://doi.org/10.1038/s41586-019-1549-9

Pyle, A., Anugrha, H., Kurzawa-Akanbi, M., Yarnall, A., Burn, D., & Hudson, G. (2016). Reduced mitochondrial DNA copy number is a biomarker of Parkinson’s disease. Neurobiology of Aging, 38, 216.e7-216.e10. https://doi.org/10.1016/j.neurobiolaging.2015.10.033

Rafelski, S. M., Viana, M. P., Zhang, Y., Chan, Y.-H. M., Thorn, K. S., Yam, P., Fung, J. C., Li, H., Costa, L. da F., & Marshall, W. F. (2012). Mitochondrial network size scaling in budding yeast. Science (New York, N.Y.), 338(6108), 822–824. https://doi.org/10.1126/science.1225720

Sagan, L. (1967). On the origin of mitosing cells. Journal of Theoretical Biology, 14(3), 225–IN6. https://doi.org/10.1016/0022-5193(67)90079-3

Shadel, G. S. (1999). Yeast as a Model for Human mtDNA Replication. The American Journal of Human Genetics, 65(5), 1230–1237. https://doi.org/10.1086/302630

Shiiba, D., Fumoto, S.-I., Miyakawa, I., & Sando, N. (1997). Isolation of giant mitochondrial nucleoids from the yeastSaccharomyces cerevisiae. Protoplasma, 198(3), 177–185. https://doi.org/10.1007/BF01287567

Simon, V. R., Karmon, S. L., & Pon, L. A. (1997). Mitochondrial inheritance: Cell cycle and actin cable dependence of polarized mitochondrial movements in Saccharomyces cerevisiae. Cell Motility, 37(3), 199–210. https://doi.org/10.1002/(SICI)1097-0169(1997)37:3<199::AID-CM2>3.0.CO;2-2

Sniegowski, P. D., Dombrowski, P. G., & Fingerman, E. (2002). Saccharomyces cerevisiae and Saccharomyces paradoxus coexist in a natural woodland site in North America and display different levels of reproductive isolation from European conspecifics. FEMS Yeast Research, 1(4), 299–306. https://doi.org/10.1111/j.1567-1364.2002.tb00048.x

Solieri, L. (2010). Mitochondrial inheritance in budding yeasts: Towards an integrated understanding. Trends in Microbiology, 18(11), 521–530. https://doi.org/10.1016/j.tim.2010.08.001

Stevens, B. (1981). Mitochondrial structure. In The molecular biology of the yeast Saccharomyces: Life cycle and inheritance (pp. 471–504). Cold Spring Harbor Laboratory Press.

Wai, T., Ao, A., Zhang, X., Cyr, D., Dufort, D., & Shoubridge, E. A. (2010). The Role of Mitochondrial DNA Copy Number in Mammalian Fertility1. Biology of Reproduction, 83(1), 52–62. https://doi.org/10.1095/biolreprod.109.080887

Westermann, B. (2014). Mitochondrial inheritance in yeast. Biochimica et Biophysica Acta (BBA) - Bioenergetics, 1837(7), 1039–1046. https://doi.org/10.1016/j.bbabio.2013.10.005

Williamson, D. H. (2002). The curious history of yeast mitochondrial DNA. Nature Reviews Genetics, 3(6), 475–481. https://doi.org/10.1038/nrg814

Williamson, D. H., & Moustacchi, E. (1971). The synthesis of mitochondrial DNA during the cell cycle in the yeast Saccharomyces cerevisiae. Biochemical and Biophysical Research Communications, 42(2), 195–201. https://doi.org/10.1016/0006-291X(71)90087-8

Williamson, D. H., & Slonimski, P. P. (1976). Packaging and Recombination of Mitochondrial DNA in Vegetatively Growing Yeast Cells. In Genetics, Biogenesis, and Bioenergetics of Mitochondria (pp. 117–124). De Gruyter. https://www.degruyter.com/document/doi/10.1515/9783111522241-012/html

Zakhartsev, M., & Reuss, M. (2018). Cell size and morphological properties of yeast Saccharomyces cerevisiae in relation to growth temperature. FEMS Yeast Research, 18(6), foy052. https://doi.org/10.1093/femsyr/foy052

Zampar, G. G., Kümmel, A., Ewald, J., Jol, S., Niebel, B., Picotti, P., Aebersold, R., Sauer, U., Zamboni, N., & Heinemann, M. (2013). Temporal system-level organization of the switch from glycolytic to gluconeogenic operation in yeast. Molecular Systems Biology, 9(1), 651. https://doi.org/10.1038/msb.2013.11

Zeviani, M., & Di Donato, S. (2004). Mitochondrial disorders. Brain, 127(10), 2153–2172. https://doi.org/10.1093/brain/awh259

