## Supplemental figures and tables for "Changes to the mtDNA copy number during yeast culture growth"

### Supplementary Figures and Tables

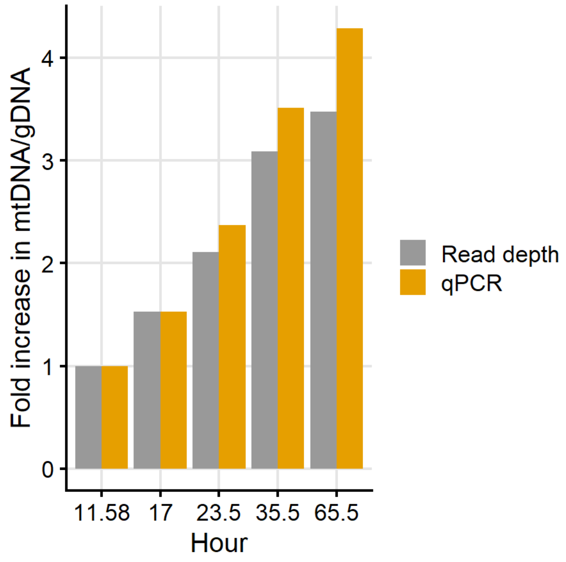


**Fig. S1.** A comparison of the two methods used to assay mtDNA fold increase during culture growth, qPCR and read depth from whole-genome sequencing. Samples were taken from the same culture at the same time. Data is same as that presented in Fig. 2B.

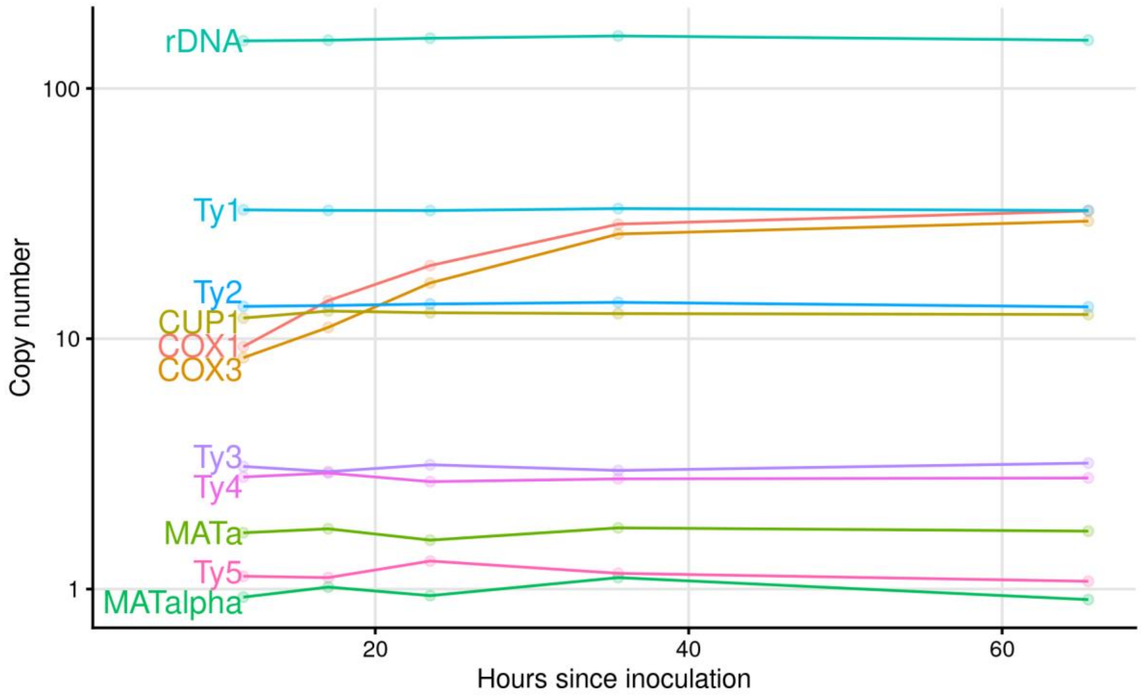


**Fig. S2.** The mitochondrial *COX1* and *COX3* genes increase in copy number during culture growth. *COX1* provides the data for mtDNA copy number in Fig 2B mtDNA fold increase in Fig. S1. As controls, we assayed the copy number of 9 genomic loci: ribosomal DNA, *CUP1*, *MATa*, *MATα*, and the 5 Ty transposons.

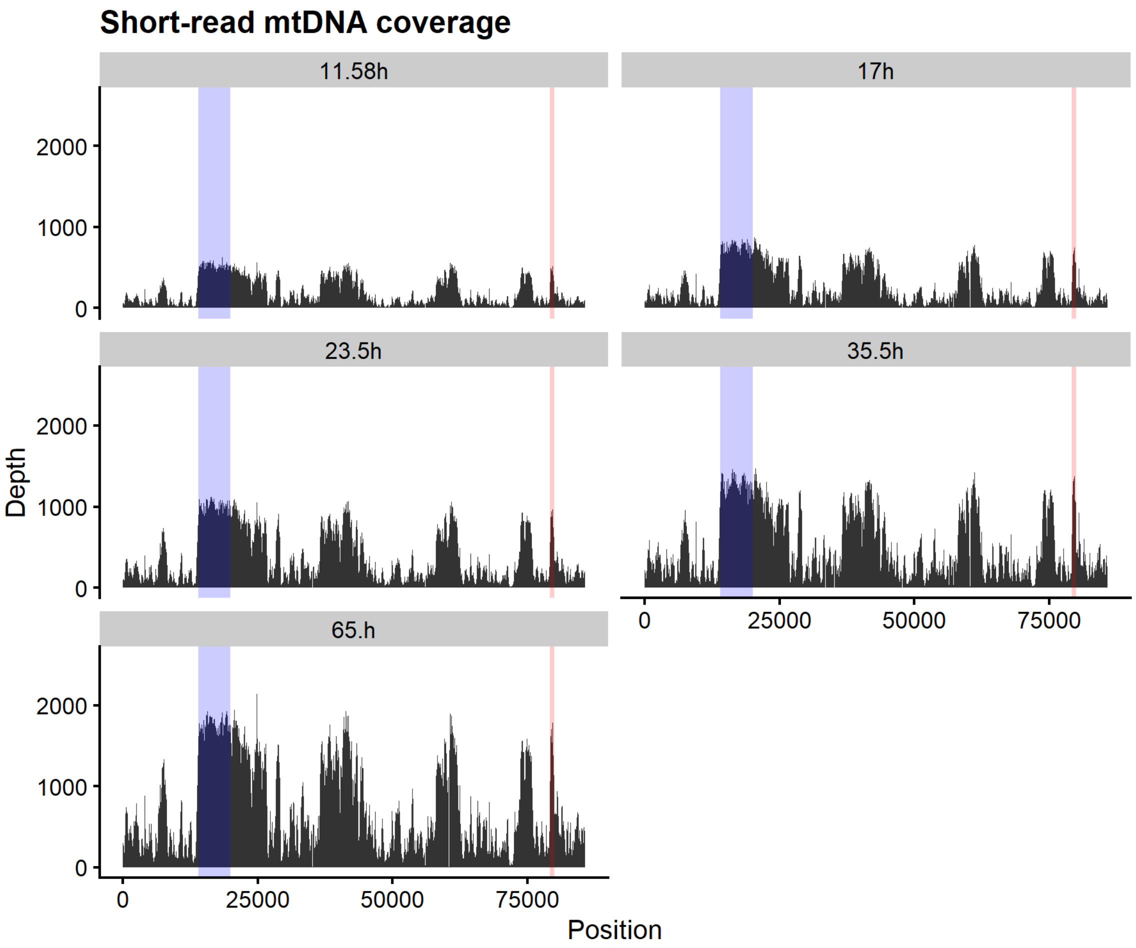


**Fig. S3.** From short-read (Illumina) sequencing, mtDNA read depth from the 5 timepoints sequenced, with regions of mtDNA used for determining copy number highlighted: *COX1* in blue; *COX3* in red.

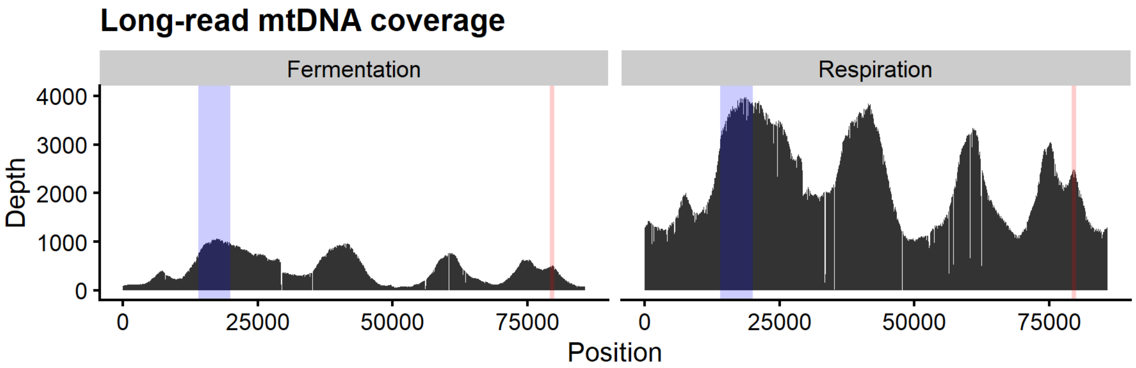


**Fig. S4.** From long-read (Nanopore) sequencing, mtDNA read depth, with regions of mtDNA used for determining copy number highlighted: *COX1* in blue; *COX3* in red.

**Table S5.** Long-read sequencing gives near identical copy number results to short-read sequencing for identical extracts. DNA extracts from the cultures described in Fig. S6 were analyzed by both short- and long-read sequencing, and the resulting mtDNA copy numbers are very similar.

| | Illumina | Nanopore |
|---|---|---|
| Fermentation | 8.6 | 8.2 |
| Respiration | 27.8 | 28.5 |

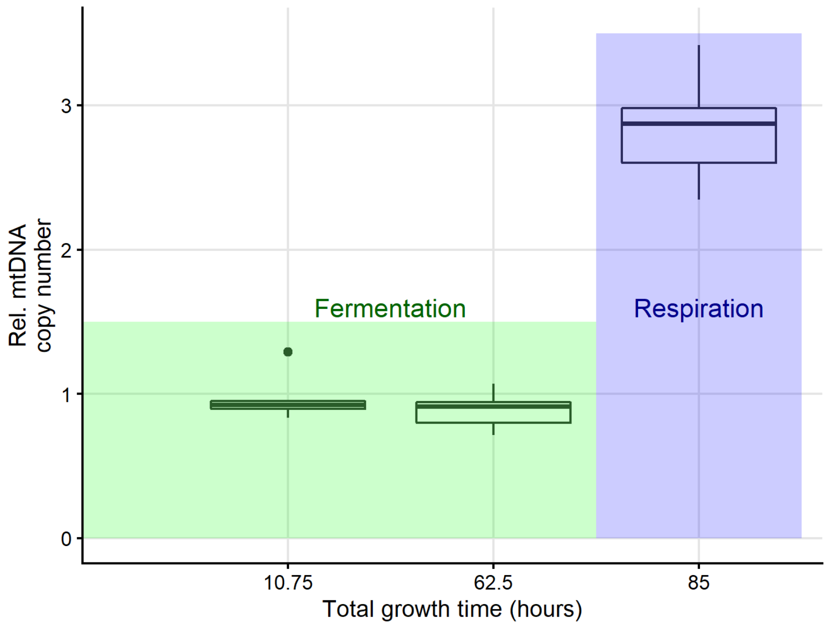


**Fig. S6.** A culture of the haploid lab strain was inoculated as described in Methods, and then three consecutive serial transfers were performed during the fermentation phase so that total consecutive time in rapid fermentative growth was over 60 hours. DNA extractions were performed during fermentation in the first culture, during fermentation in the fourth culture, and during respiration in the fourth culture. This experiment demonstrates that even after a very long period of time in fermentation, the mtDNA copy number remains low.

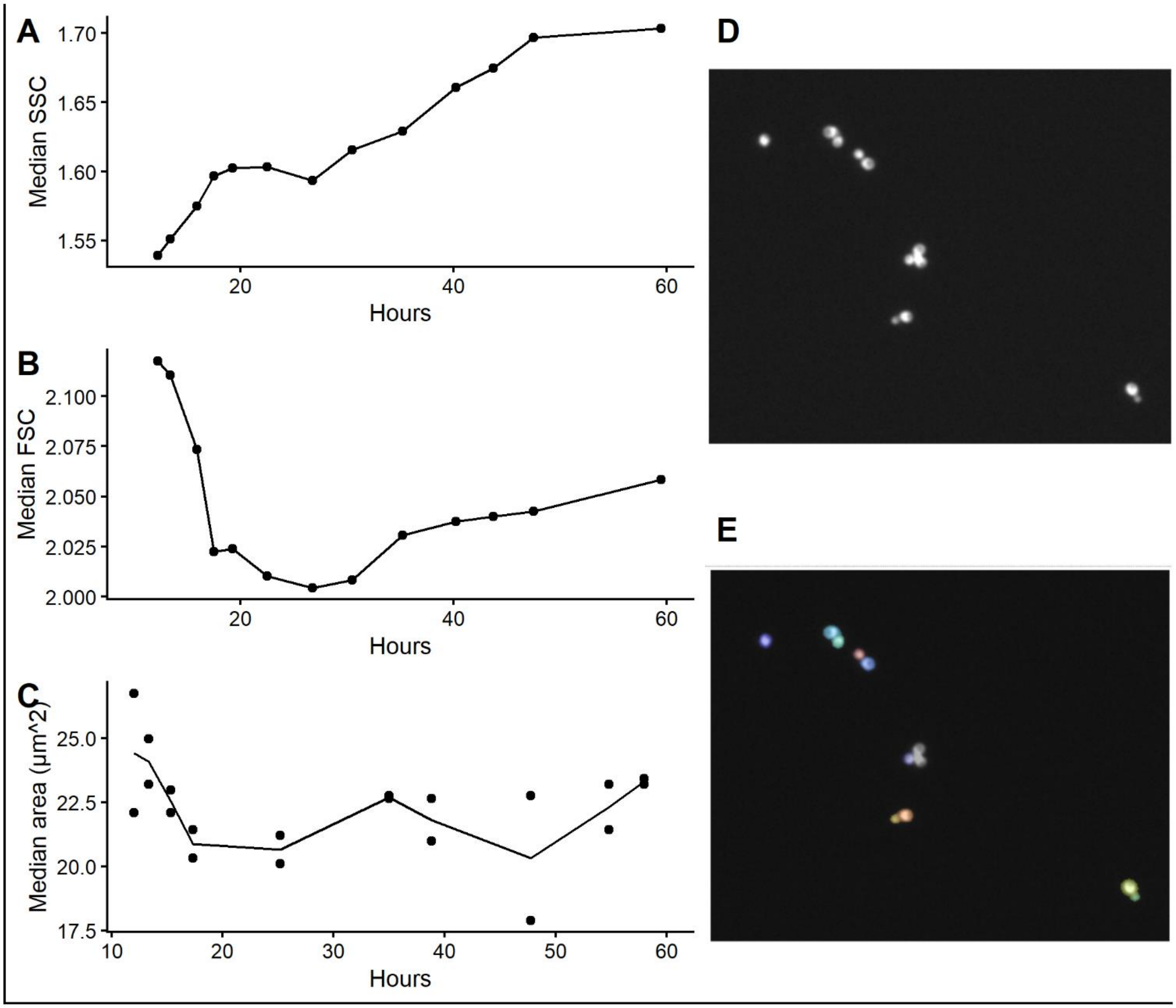


**Fig. S7. (A)** Median side scatter as a function of hours after inoculation, measured for the experiment shown in Figure 2 (a haploid W303 strain). **(B)** Median forward scatter (FSC), a proxy for cell size. **(C)** A microscopy-based live imaging assay from a similarly grown culture. **(D)** Sample image and subsequent object detection **(E)** for the imaging assay. The gray triplet was not declustered accurately so it was removed manually.

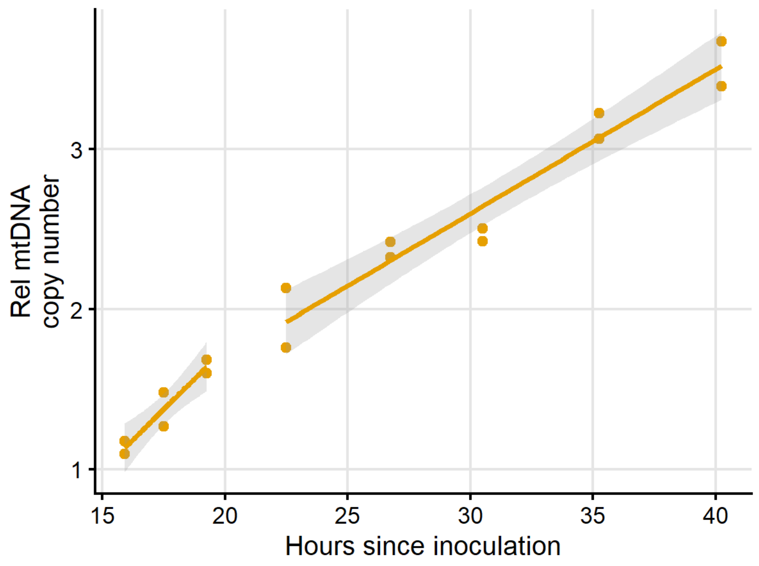


**Fig. S8.** This plots the same data as Fig 2A. The slope of the mtDNA increase during the diauxic shift (0.15 units/hr) is greater than after (0.09 units/hr). The difference is not statistically significant (no significant interaction term in a linear regression, $p$=0.40).

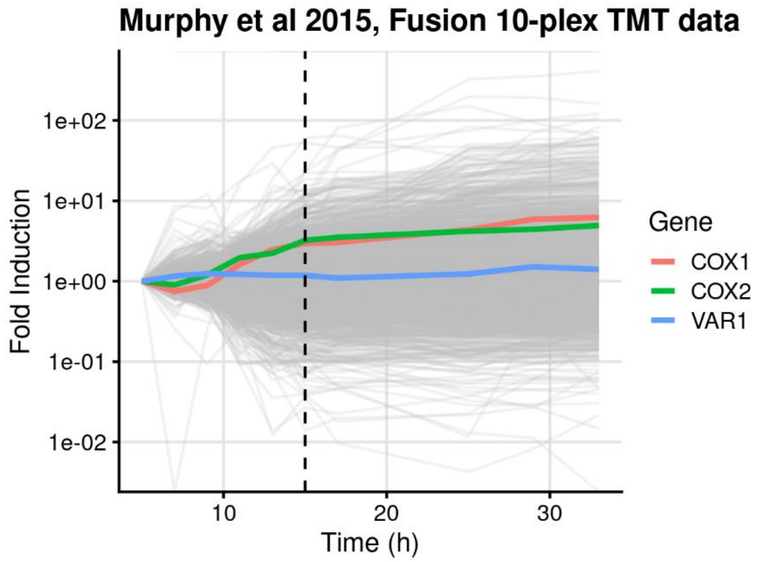


**Fig. S9.** Re-plotted from proteome data provided by Murphy et al. 2015 with three genes located on the mitochondrial chromosome highlighted. The dashed line represents glucose depletion. *COX2* was characterized by the authors as being in the group of genes that are "early induced" with respect to the diauxic shift. *COX1* and *COX2* encode proteins of the core cytochrome *c* oxidase complex. *VAR1* encodes a mitochondrial 37S ribosomal protein.
